# Machine learning models of human tissue microbiomes for tissue-of-origin prediction

**DOI:** 10.1101/2024.05.06.592823

**Authors:** Gita Mahmoudabadi, Stephen R. Quake

**Affiliations:** Department of Bioengineering, Stanford University, Stanford CA, USA; Department of Applied Physics, Stanford University, Stanford CA, USA; Chan Zuckerberg Initiative, Redwood City, CA, USA

## Abstract

There is increasing interest in using microbial data diagnostically for tissue health monitoring such as in early cancer detection. To build such models, we need to understand whether normal tissue microbiomes can also be predictive of tissue of origin, and importantly ask how contaminants may contribute to model performance. In this study, using the Tabula Sapiens Microbiome dataset, we built machine learning models of human tissue microbiomes that can predict tissue of origin. This may in part explain how tumor types can be predicted based on the tumor microbiomes. We also demonstrate that machine learning models built using contaminants alone, though not as powerful as those built on true signal, can still predict tissue of origin. Reassuringly, the addition of contaminants to true signal does not increase the performance over models built on true signal. Overall, our findings raise the burden of proof for predictive models of the human tissue and tumor microbiomes. Without addressing the magnitude of contribution from contaminants to model performance, a model’s reproducibility and its clinical value becomes questionable. We also discuss the optimal microbial taxonomic resolution for building these models.

## Introduction

Recent advancements in human microbiome research have provided profound insights into the role of microbial communities across a range of metabolic, neurological, cardiovascular, and autoimmune diseases^1–6^. Traditionally, the focus of microbiome studies has been on accessible sites (or what we refer to as external-facing microbiomes) such as the gut, oral cavity, and skin, with internal sites presumed largely sterile. However, emerging research has challenged this assumption, and has uncovered microbial signatures within human tumors and healthy tissues. For example, recent studies have identified tumor-specific mycobiomes and microbiomes^7,8^. Other studies have further demonstrated the existence of microbes within tumors and in some instances shown that the microbiome can contribute to tumorigenesis and tumor progression^9–17^.

Since the discovery of microbial and fungal reads in tissues and tumors, there is growing interest in building predictive models for cancer diagnostics and prognostics using this data^7,18–21^. However, systematic pan-tissue surveys of normal tissue microbiomes are still largely lacking, and yet they are needed for interpreting models built for tumor detection. For example, if one can build models that can predict tissue of origin from microbial data derived from normal tissues, to what extent are models built on tumor microbiomes truly specific to tumors as compared to surrounding normal tissue? Additionally, tissues and tumors represent samples with very low microbial biomass, and in such settings, the impact of contaminants can be substantial. Thus, another important largely missing facet so far is the quantification of the contribution of contaminants to the performance of machine learning models. In this study, we aimed to address these two topics.

We previously discovered human tissue and cell type microbiomes by using the raw

sequences from the Tabula Sapiens^22^ which encompasses 400,000+ annotated human cells, encompassing 100+ cell types and 19 tissues from 11 human organ donors (**Figure 1**). Through highly stringent experimental and computational decontamination and quality control filters, we identified viral, bacterial, and fungal sequences originating from human single-cell transcriptomes, together forming the Tabula Sapiens Microbiome (TSM) dataset^23^. By integrating the TSM with 20+ external, large-scale datasets, we were able to benchmark the precision of our computational approach and map out the possible flow routes of microbes from external-facing microbiomes to internal tissues and tumors. We showed that tumor microbiomes share many common taxa with tissue microbiomes extracted from normal tissues. We further validated our findings based on single-cell sequencing using DNA sequencing and imaging on tissues from a separate cohort of 8 donors. Finally, we shed light on the types of organisms that human cells encounter by physical proximity to microbial niches and sites of infections, or through immune clearance, or in more rare cases, by hosting an intracellular organism such as Epstein Barr Virus (EBV).

**Figure 1.**
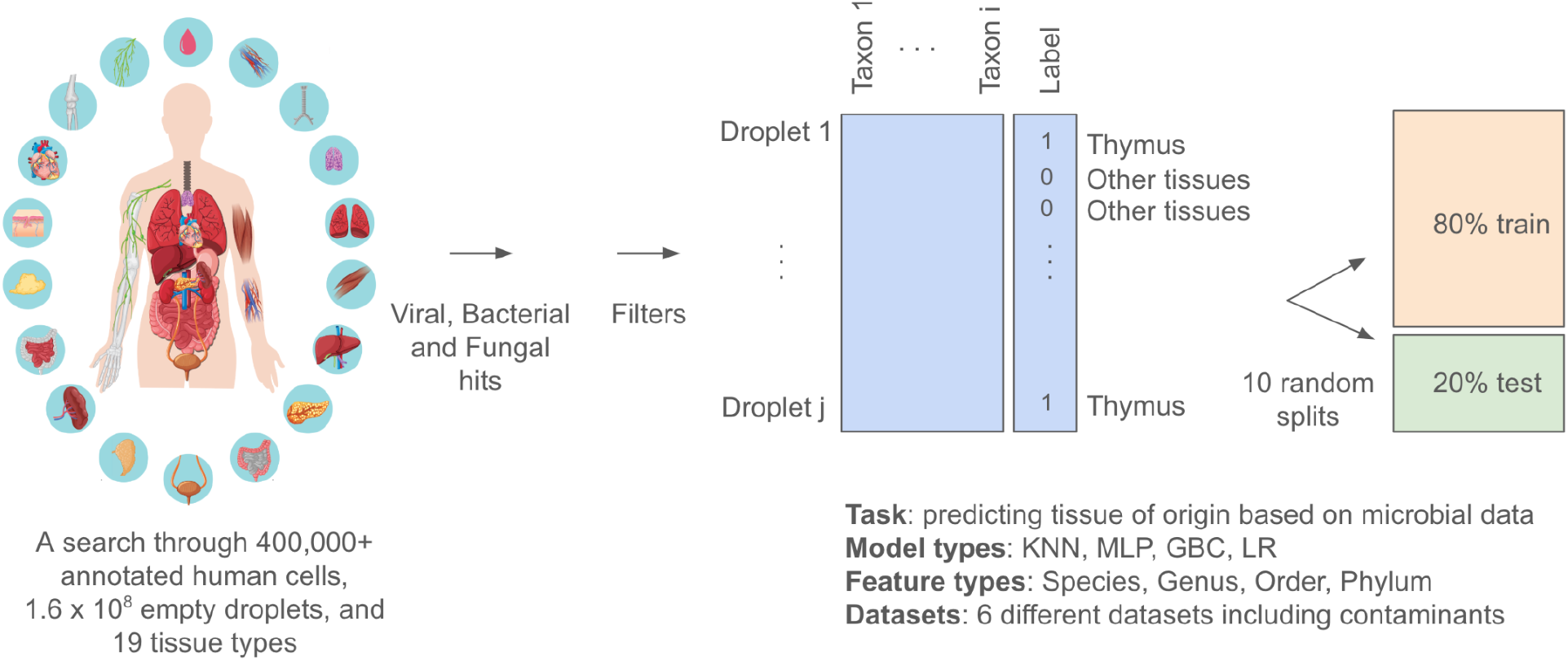
Study design. The Tabula Sapiens Microbiome (TSM) dataset derived from searching through human tissue derived droplets (empty or containing cells) as previously described by Mahmoudabadi et al.^23^ In this study various feature tables (shown in blue) are constructed where the droplets are rows and taxa are columns. The table holds the relative abundance of each taxa in each droplet. For each feature table, ten models are built using ten random train and test splits. The model types included in this study are K-Nearest Neighbors (KNN), Multi-layer Perceptron (MLP), Gradient Boosting Classifier (GBC) and Logistic Regression (LR). The taxa types tested include species, genera, orders, and phyla. Datasets included in this study are shown in SI Figure 1.

## Results

### Models built on genus- or species-level taxa provide the best tissue-of-origin predictions

Having built the TSM dataset, we wondered whether we could build machine learning models of human cellular and cell-free microbiomes that could be predictive of the tissue of origin. A prior study used machine learning models to identify tumor types based on the tumor microbiomes, suggesting tumor microbiomes can serve as a potential diagnostic signal^24^. However, an important limitation of this study is that it lacked the negative control samples associated with the experimental protocol. Similar studies which have experimentally accounted for contaminants also have not quantified to what extent contaminants can provide predictive signal and potentially boost model performances if they are missed or undersampled^7,18,25^.

As detailed in the previous study, we devoted five 10X lanes to deeply sequence ∼200 negative control samples derived from reagents and instruments used during the experimental protocols^23^. While these samples were collected post tissue processing experiments, they were acquired from the same batch or from the same exact sample used in the experiments that were then stored in freezers. Eliminating taxa found in the contamination dataset constituted the first filter on the dataset. The portion of the dataset that does not pass through this filter is referred to as the contamination dataset. To eliminate any potentially missed contaminants we also applied two filters based on commonly found bacterial and fungal contaminants reported in previous studies. We then built machine learning models based on known and putative contaminant signatures to identify additional putative contaminants in the TSM dataset (Filter 4). Subsequent filters (Filters 5-7) were applied to select for longer alignment lengths, and focus the resulting hits on microbial taxa that have been observed in previous studies as part of the external-facing microbiomes. The dataset that has gone through all these seven filters is referred to as TSM F7, or F7 for short. To gauge the potential contribution of contaminants to model performance, in this study we compare machine learning models of the human tissue microbiomes to those built using contaminants alone.

To build machine learning models of the human tissue microbiome using the TSM F7 dataset, we created a feature table where the rows are droplets and the columns are microbial taxa (see Methods). **SI Figure 1** displays the datasets used in this study. For each model, we dedicated 80% of the data for training and 20% for testing. We generated one-tissue-versus-rest models, using the thymus as the tissue of interest since it is one of the tissue with the most number of microbial hits in the TSM dataset. We tested the performance of several ML model types, namely logistic regression (LR), gradient boosting classifier (GBC), k-nearest neighbor (KNN), and multi-layer perceptron (MLP) on this feature table at selecting a tissue from all the rest. For each model type, we built ten models using ten random train/test splits and report the resulting Area Under the Precision-Recall (AUPR) and Area Under the Receiver Operator Curve (AUROC) values (**Figure 2**). Because AUPR is often the relevant metric when evaluating imbalanced datasets such as those used in one-versus-rest models, we generally rely on this metric to make selections. We found the GBC model type provides the best AUPR confidence interval, however, the slightly better performance of the GBC model type is not statistically significant compared to other model types (Mann-Whitney one-side U tests, N=10 per group, P values<0.05), thus other model types can also be used to produce similar results.

**Figure 2.**
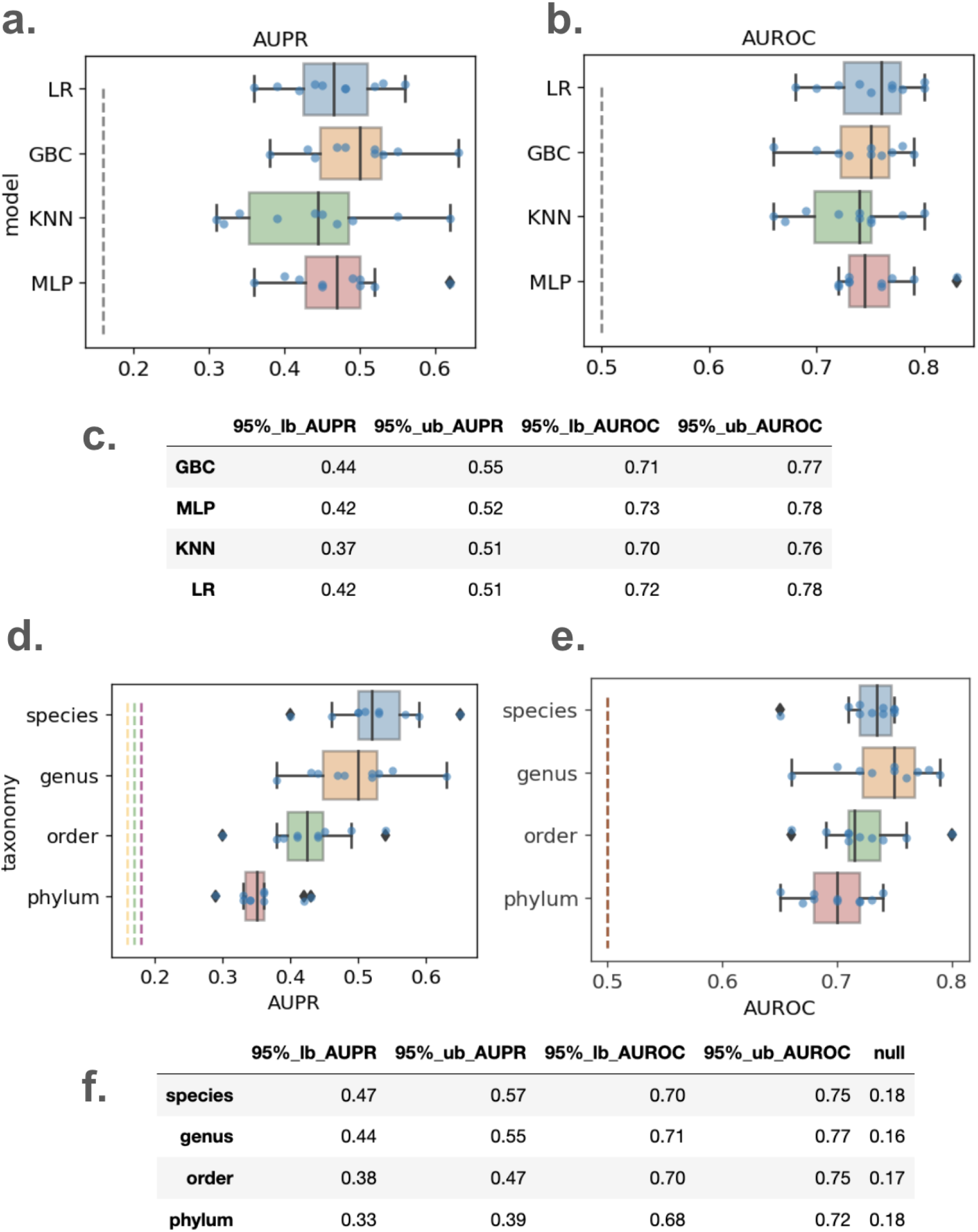
Model AUPR and AUROC performances for thymus-versus-rest prediction task across different model types a-c (LR: logistic regression, GBC: gradient boosting classifier, KNN: k-nearest neighbor, MLP: multi-layer perceptron) and across different taxonomic resolution (d-f) applied onto TSM genera withheld dataset. Results from 10 separate iterations of each model type are shown (10 different train-test splits). The null AUPR and AUROC values are shown by the dashed gray lines. The 95% confidence intervals are shown in tables (c and f) (lb: lower-bound, ub: upper bound).

We proceeded to test the performance of the GBC thymus-verus-rest model when given four different feature tables constructed with species, genera, orders and phyla as features. We found that models trained with species or genus-level feature tables provide best performances (**Figure 2**). Models trained on order or phylum level features perform significantly worse (Mann-Whitney one-side U test on AUPR values comparing order to genus level tables, n=10 for each group, U=24, P<0.05). This is simply because as taxonomic resolution is decreased there is an important differentiating signal that is lost. As we have shown before^23^, genus-level taxonomic resolution is optimal for precision and recall of short read (∼100 bp) profiling. And because there is no significant performance difference between species-level and genus-level models (Mann-Whitney one-side U test on AUPR values, n=10 for each group, U=62, P>0.05) while genus-level models are more computationally efficient, we chose to build subsequent models using genus-level classification.

### Models based on contaminants can provide tissue-of-origin prediction but perform worse than models based on true signal

Next, we compared the performance of the GBC model against the TSM F7 and the contamination datasets, using genus-level categorization to construct the feature table. This dataset is 1686 (number of droplets) by 107 (number of genera). As with many other examples in machine learning, model performance is highly impacted by the size of the training data. Thus, to compare between models constructed using the contamination and the F7 datasets, we downsampled the contamination dataset to the same number of droplets as the F7 dataset, so that it is a matrix of 1686 by 416. Additionally we matched it to the F7 dataset in terms of its prevalence of positive class (i.e. proportion of droplets from the thymus compared to other tissues). We show that models that are given F7 data perform better than those given a matched contaminant dataset (Mann-Whitney one sided U test, P-value < 0.05) despite a greater number of taxa appearing in the contaminant dataset (**Figure 3.a**).

**Figure 3.**
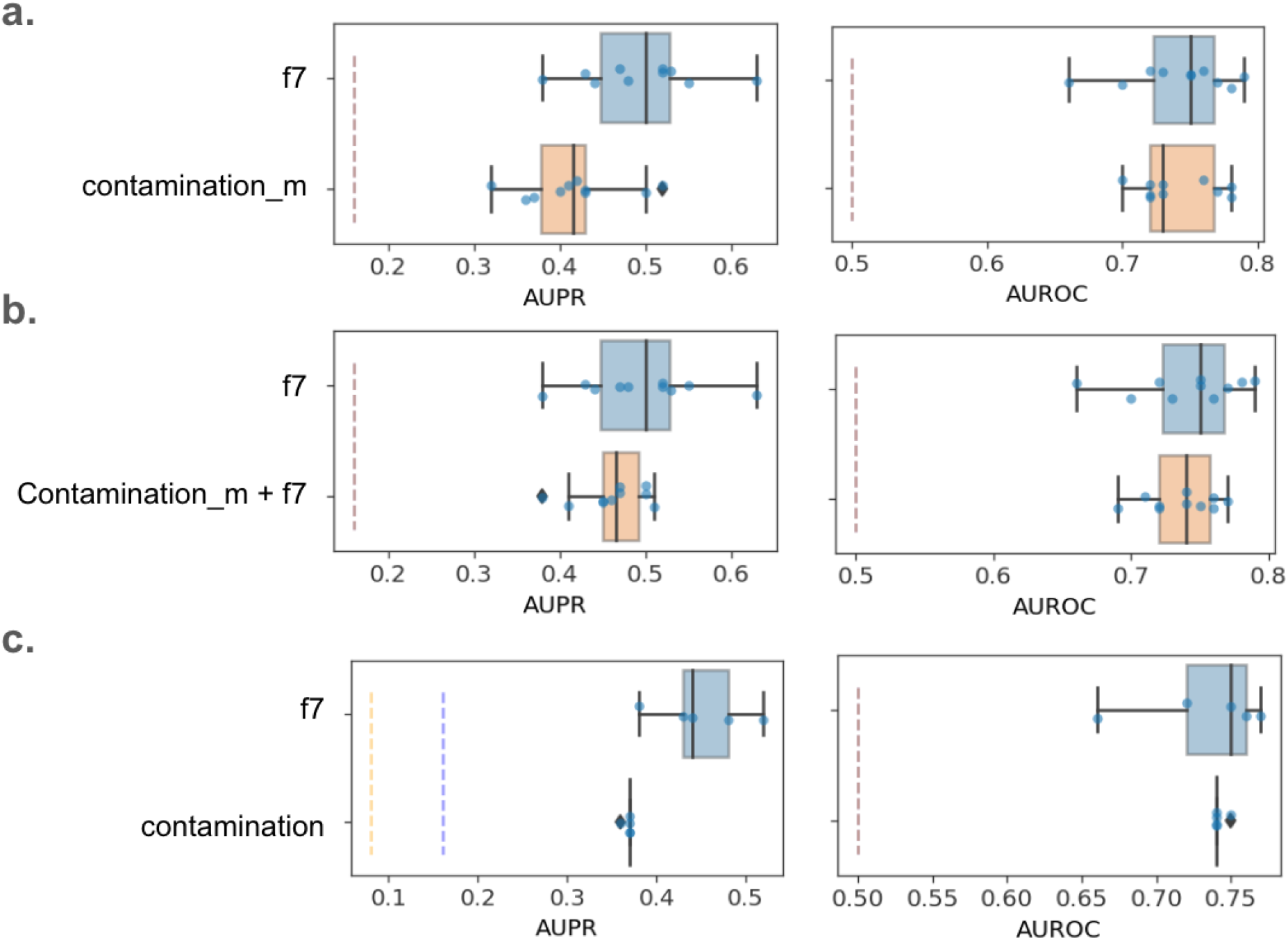
Area under the precision-recall curve (AUPR) and area under the receiver-operator curve (AUROC) for one-tissue-versus-rest Gradient Boosting Classifier (GBC) applied onto various datasets. a) comparison of models built on the f7 dataset to those built on the matched contamination dataset. b) comparison of models built on the f7 dataset to those built on a combined dataset of f7 and matched contamination. c) comparison of models built on the f7 dataset to those built on the full contamination dataset. 10 separate iterations of the GBC model are shown, with different train-test splits. The null AUPR values(prevalence of the positive class) are shown as dashed lines (blue: F7). The null AUROC value is 0.5.

### Models based on a mixture of contaminants and true signal do not outperform models built on true signal

Given that the F7 and the contamination based models can be predictive of tissue of origin, albeit to different extents, we wondered whether in a situation where contaminants are unknown and cannot be subtracted out, how they might impact the resulting models. To test this scenario, we concatenated the two datasets (F7 and matched contamination), such that the mixed dataset had both greater number of droplets and genera for training and testing (matrix of 3372 x 496). Surprisingly, the addition of the two datasets did not increase the performance over models built on the F7 dataset. These models though do not perform worse than models based on F7 alone (Mann-Whitney one sided U test on AUPR, U=32.5, N=10 in each group, P-value > 0.05; Mann-Whitney one sided U test on AUROC, U=41.5, N=10 in each group, P-value > 0.05) (**Figure 3.b**). Hence, there is an interesting non-synergistic effect at play where there is no added benefit to the model performance when there is additional data from contaminants, even though on their own contaminants have predictive value.

When the F7 and the contaminant datasets are not matched (matrix of 445,241 x 416), the models trained on contaminants still do not outperform despite the vast increase in the training dataset size (**Figure 3.c**). They produce models with statistically lower AUPR values (Mann-Whitney one sided U test, U=0, N=10 in each group, P-value < 0.05), and similar AUROC values (Mann-Whitney one sided U test, U=41, N=10 in each group, P-value > 0.05).

### Comparison of one-versus-all models for more than a dozen tissues using true signal and contaminants

We proceeded to create one-versus-all GBC models for 15 tissues in TSM with the most number of microbial hits. We report the AUROC and AUPR results in **SI Figure 2**. Most tissues contain enough microbial data to enable models with AUPR and AUROC values above the null, however, with a larger dataset these values will likely substantially improve. The reason for this assumption is that to increase the number of microbial observations per tissue, we relaxed our previous stringent filters and included hits from all genera that had appeared in the F7 dataset but still eliminated species found in the contamination dataset. We call this dataset, the F7 expanded dataset, or F7_e for short. It is still stringent in the sense that it includes members of genera that have passed through all filters, but it provides roughly 4 times the number of microbial observations (or hits) as the F7 dataset. The one-versus-all models on the F7_e dataset are shown in **Figure 4**.

**Figure 4.**
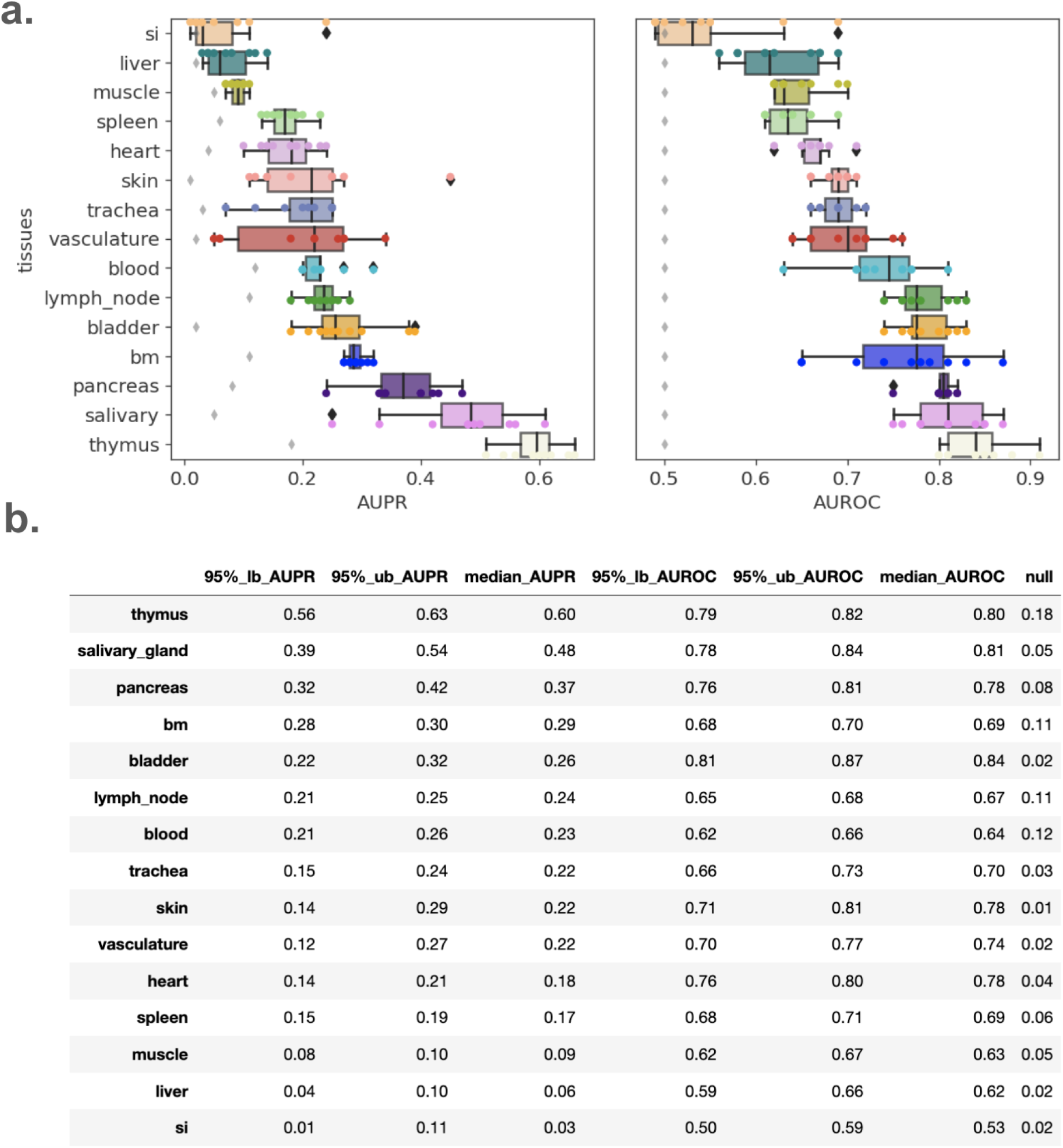
Area under the precision-recall curve (AUPR) and area under the receiver-operator curve (AUROC) for one-tissue-versus-rest Gradient Boosting Classifier (GBC) applied onto the f7_e dataset. a) 10 separate iterations of the GBC model are shown, with different train-test splits. The diamonds indicate null performance. The plots are ordered by median values. b) The 95% AUPR and AUROC confidence intervals along with median values are shown for each tissue (bottom table).

To compare the performance of the one-versus-all models using the F7_e and F7 datasets, we identified the top five tissues with the highest median AUPR values for each dataset. For the F7_e models the list of top tissues includes thymus, salivary gland, pancreas, bone marrow and bladder. For the F7 dataset, the list is thymus, bladder, lung, lymph node and spleen. We compared one-versus-rest models for these tissues for a total of five comparisons: Mann-Whitney one-sided U tests, U=[13, 35.5, 47.5, 12.5, 27.5], P values=[<0.05, >0.05, >0.05, <0.05, <0.05]. For the two non-significant comparisons, we tested to see if F7 models were significantly better, and we see that they are not (Mann-Whitney one-sided U test, U=[64.5, 47.5], P=[>0.05, >0.05]). Thus, we conclude that as expected, by supplementing models with additional data, they generally perform better.

We then repeated the same exercise to compare the F7_e models to the contaminant dataset matched to the size of the F7_e dataset in terms of the total number of droplets (**SI Figure 3**). The resulting U values were [38, 11.5, 1, 7, 18] and the resulting P values were [>0.05, <0.05, <0.05, <0.05, <0.05]. Hence, as we have seen with previous comparisons of F7 to the full contamination dataset, models based on contaminants do not perform as well.

## Discussion

In this study we showed that given microbial data extracted from a tissue, the human cellular and cell-free microbiomes can be predictive of the tissue of origin. Under various settings we showed that the models built on true signal perform better than models built on the contamination dataset. However, this quantifies an important issue in this field, namely that models based on contaminants alone can also learn tissue of origin. Why does contamination enable predictive models? The reason is likely that much of the reagents used for tissue processing experiments are tissue-specific.

With increasing interest in building predictive models of tissue and organ health based on microbiomes, it is imperative that there is comparison of these models to datasets of known contaminants physically captured from every aspect of the experimental protocol. Without a careful consideration of how a model performs when given pure contamination, it would not be clear how much of the model’s performance is attributable to true microbial signal.

With careful controls, increasingly powerful machine learning models combined with increasingly large datasets for training could begin to guide us on the impact of the tissue microbiome on tumor origins, progression and escape. They can potentially be diagnostic of other important human diseases, particularly autoimmune, metabolic and neurological diseases through microbial interactions with the human immune, digestive and nervous systems.

## Limitations

While the models shown in this study have predictive power, there is much room for improvement. In building the TSM F7 dataset, because it represented one of the first attempts at a holistic map of microbial sequences in human tissues using single-cell data, our goal was to prioritize precision of microbial identification over recall to mitigate false positives as much as possible. Additionally, single-cell transcriptomics is not optimized for capturing microbial sequences which often lack poly-adenylated sequences. Finally, we were tasked with high-precision detection of microbial sequences in the background of sequences that are predominantly human in origin. Together all these challenges limit the number of microbial sequences that we are able to capture and use to train models on. However, with methodological improvements, there is an incredible opportunity for further investigating the human tissue microbiome and its impact on various areas of human health. The machine learning models’ performances will also likely increase substantially with more training data.

## Conclusion

We previously assembled human tissue and cell type microbiomes by mining 400,000+ annotated human cells and many more cell-free droplets that encompass microbial signals from numerous human cell types and 19 tissues from 11 human organ donors. Using this Tabula Sapiens Microbiome dataset, here we showed that machine learning models can predict tissue of origin using microbial data extracted from different tissue types. However, what we also demonstrate in this study is that machine learning models can be nearly as powerful in predicting tissue of origin from contaminants alone. This is most likely because the reagents and instruments used for tissue (and tumor) processing are tissue-specific. Thus as we use increasingly powerful models to leverage the microbiome as a diagnostic frontier, it is imperative that we continue to quantify and directly model the contribution of contaminants to ensure the reproducibility of the resulting models across different settings.

## Materials and Methods

The input to machine learning models of the human tissue microbiome were feature tables where the rows are human cells and the columns are microbial genera (though we did also test species, orders and phyla as columns). This table holds the counts of microbial genera normalized by the total counts in each row. We tested the performance of several ML model types (logistic regression, gradient boosting classifier, k-nearest neighbor, and multi-layer perceptron) on this feature table. We chose thymus as the tissue to build one-versus-rest models on because it is the tissue with the most microbial hits. Thus for each row, the labels that these models were trained to predict were “thymus” (label =1) and “other” (label=0), where “thymus” is the minority, positive class.

To compare the statistical significance of model performances, we use 10 different train-test splits to build different models and report only performance on the unseen portion of the data. Using the data from 10 iterations, we report the 95% confidence intervals for AUPR and AUROC. We use the Mann-Whitney U test throughout to avoid any assumptions about the normality of the underlying data.

To match the contamination and F7 datasets, we downsampled the contamination dataset to the F7 dataset both in the number of droplets positive for hits, as well as the ratio of thymus-to-rest droplets, which represent the prevalence of the positive class, or the null AUPR. We generated 10 different contamination-matched datasets and built 10 different thymus-verus-rest GBC models.

## Code and Data Availability

Please see our GitHub repository: https://github.com/gitamahm/machine_learning_models_human_tissue_microbiomes

## Acknowledgements

We thank members of the Quake lab and the Tabula Sapiens consortium members, especially Bob Jones and Sheela Crasta. We are grateful to Donor Network West personnel including Ahmad Salehi and Ravi Ponnusamy. This work was supported by the Chan Zuckerberg Biohub, the Gordon and Betty Moore Foundation and the John Templeton Foundation.

## Author contributions

Conceptualization: GM, SRQ

Methodology: GM, SRQ

Investigation: GM, SRQ

Data Curation: GM

Formal Analysis: GM

Visualization: GM

Software: GM

Funding acquisition: GM

SRQ Resources: GM, SRQ

Project administration: GM, SRQ

Writing – original draft: GM

Writing – review & editing: GM, SRQ

## SI Figures

**SI Figure 1.**
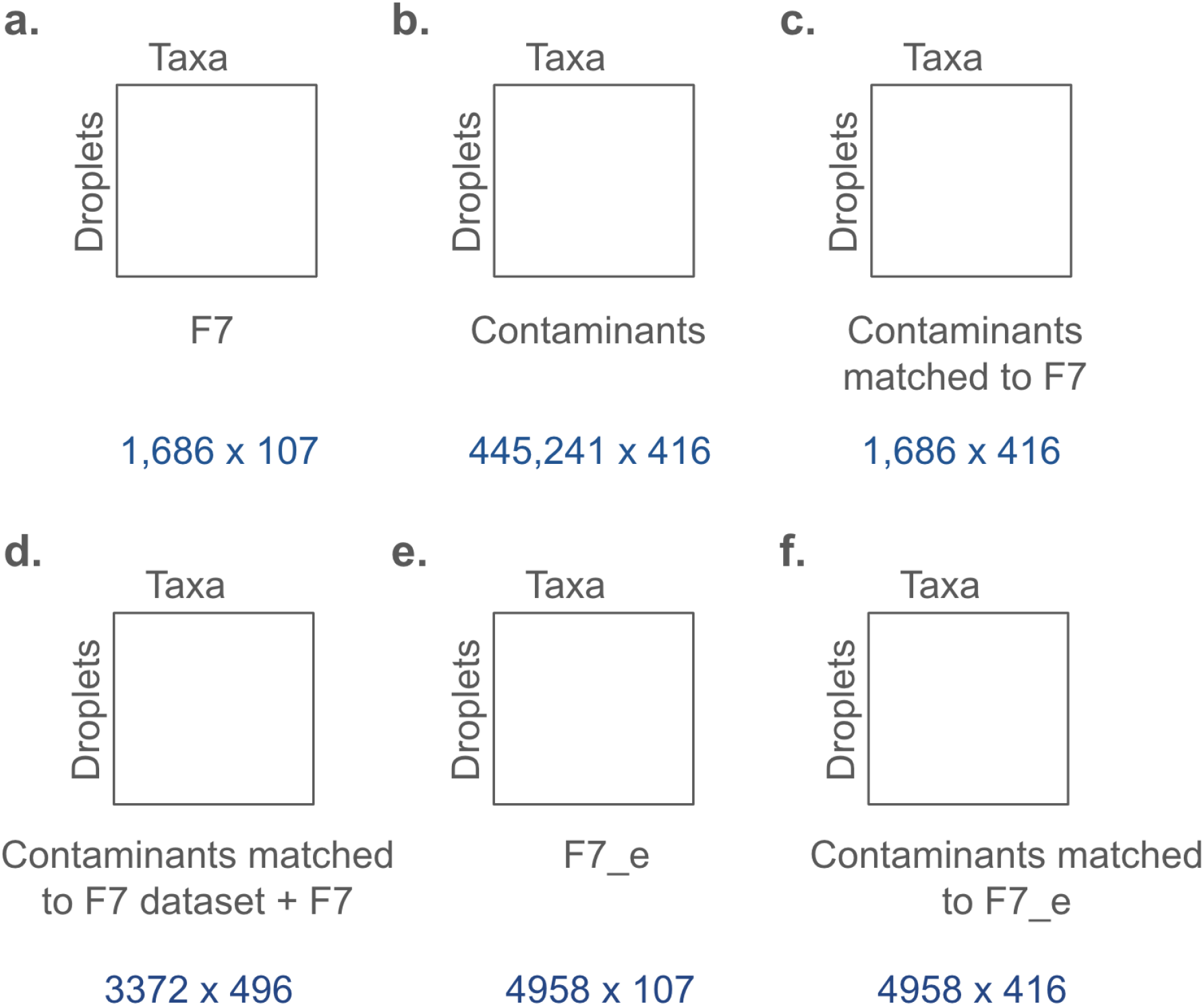
Feature tables used in this study. The feature tables are droplets as columns and taxa as rows. The size of each dataset is shown in blue.

**SI Figure 2.**
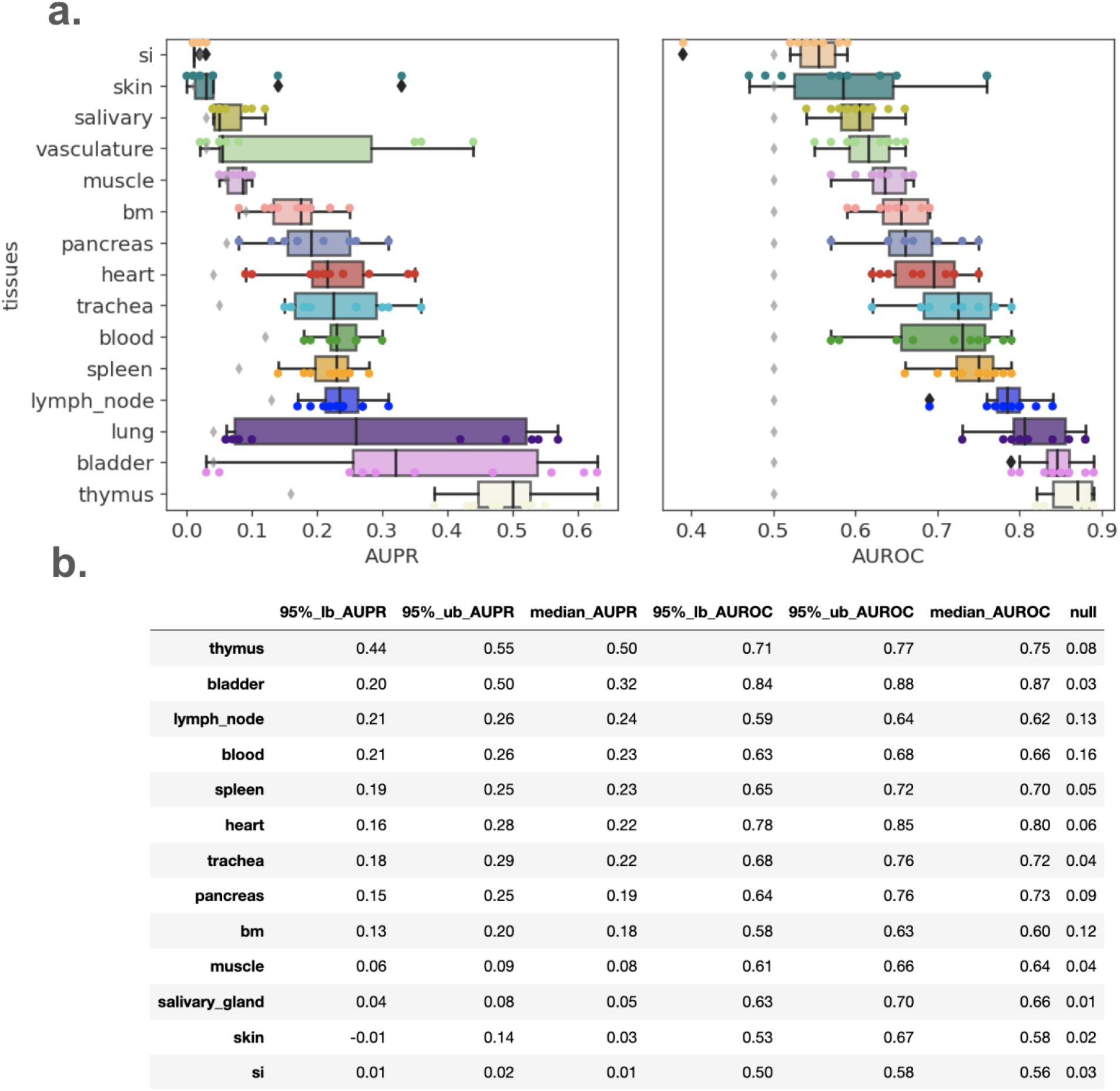
Area under the precision-recall curve (AUPR) and area under the receiver-operator curve (AUROC) for one-tissue-versus-rest Gradient Boosting Classifier (GBC) applied onto the f7 dataset. a) 10 separate iterations of the GBC model are shown, with different train-test splits. The diamonds indicate null performance. The plots are ordered by median values. b) The 95% AUPR and AUROC confidence intervals along with median values are shown for each tissue (bottom table).

**SI Figure 3.**
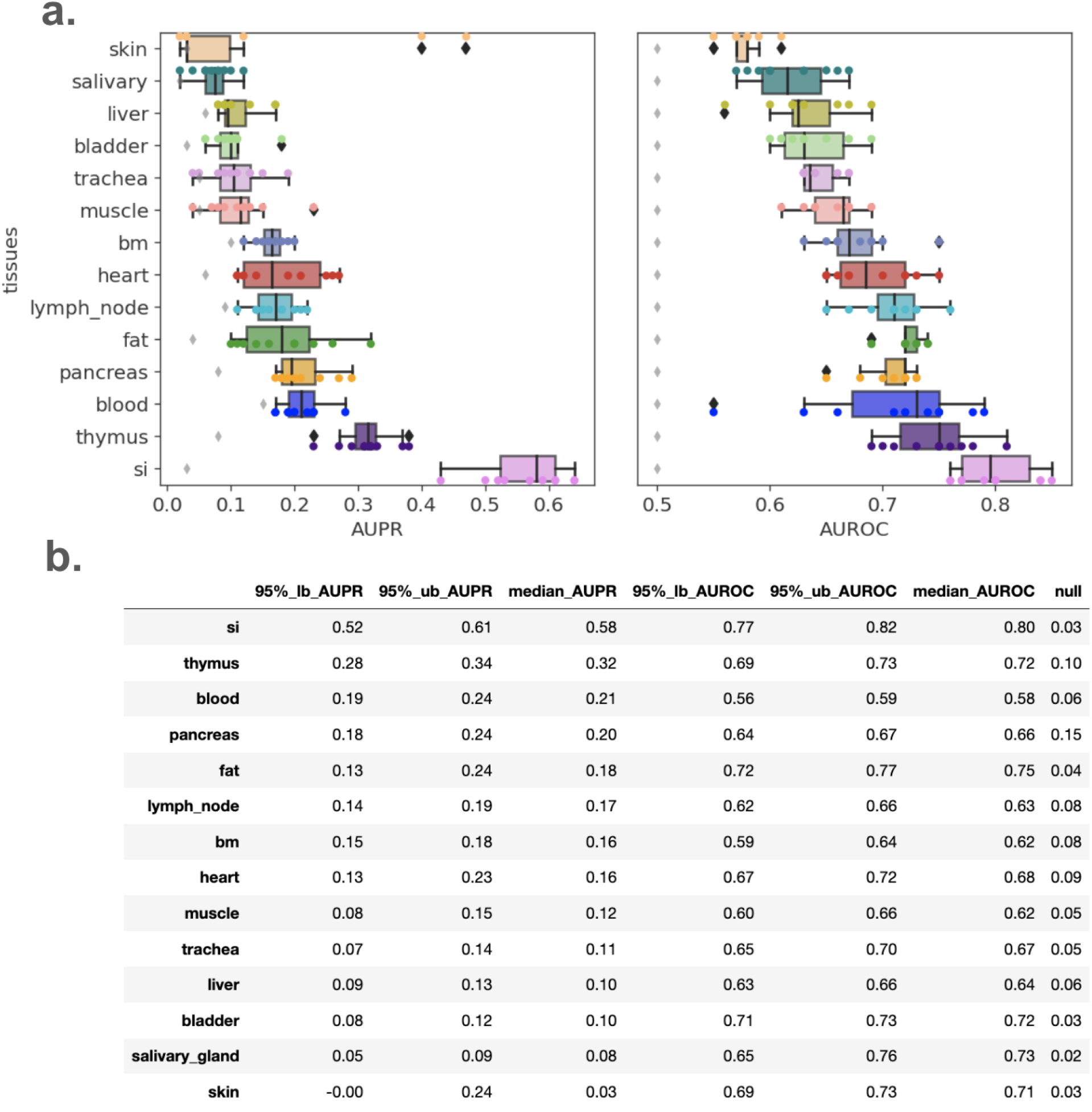
Area under the precision-recall curve (AUPR) and area under the receiver-operator curve (AUROC) for one-tissue-versus-rest Gradient Boosting Classifier (GBC) applied onto the contamination dataset matched to the f7_e dataset in terms of the number of droplets. a) 10 separate iterations of the GBC model are shown, with different train-test splits. The diamonds indicate null performance. The plots are ordered by median values. b) The 95% AUPR and AUROC confidence intervals along with median values are shown for each tissue (bottom table).

